# Macroinvertebrate diversity patterns in a guano-rich temperate cave

**DOI:** 10.1101/2025.02.09.637321

**Authors:** Luisa Dainelli, Alejandro Martínez, Fabrizio Serena, Leandro Gammuto, Caio Graco-Roza, Joachim Langeneck, Stefano Mammola, Giulio Petroni

**Author notes:** shared last author (Joachim Langeneck, Stefano Mammola, Giulio Petroni).

## Abstract

Most caves in temperate regions are energy-poor, leading to low richness and abundances of terrestrial macroinvertebrates. However, caves with abundant bat populations may represent a notable exception, as bat guano serves as a significant source of organic matter, supporting diverse specialized invertebrates. Yet, given the overall rarity of guano-rich caves in temperate region, their ecological dynamics are poorly understood. We conducted a year-long monthly standardized monitoring of the macroinvertebrates of the Buca dei Ladri cave, a guano-rich cavity in Tuscany, Italy. At each survey, we counted species within different sampling quadrats and collected environmental variables such as temperature, structural heterogeneity of the substrate, dominant substrate, and presence of bats. Over the year, we identified 31 macroinvertebrate species in the cave as both morphospecies and molecular lineages, based on mitochondrial COI and nuclear 18S genes. The latter approach hinted at the presence of cryptic diversity in some species. We observed an increase in species diversity with increasing structural complexity and the presence of bats, whereas a lower species diversity was found in guano-dominated substrates, probably due to the old age and subsequent low nutritional value of the historical guano deposits. Changes in the abundance of species within the cave were primarily explained by differences in structural complexity across sampling quadrats, whereas the distance from the cave entrance mostly explained species replacement. Our results underscore the importance of preserving healthy bat populations and a high habitat heterogeneity within the cave for the long-term conservation of the unique macroinvertebrate assemblages of the cave.

## Introduction

Terrestrial subterranean environments, such as caves, mines, and aquifers, share distinct abiotic conditions like complete darkness and stable temperatures in the deepest zones (Badino 2010; Mammola 2019). The absence of light is a critical selective factor shaping subterranean communities (Culver and Pipan 2016). Without light, photosynthesis, which forms the basis of most surface food webs, is precluded (Gibert and Deharveng 2002). Consequently, subterranean food webs rely almost exclusively on external organic resources brought by wind, rainfall, or biological vectors (Pellegrini and Ferreira 2013; Simon 2019). A significant biological vector is guano deposition (i.e., fecal accumulation by bats and other flying organisms; see Ferreira 2019). Guano supplies caves with organic matter, driving the trophic network of many invertebrate communities (Ferreira 2019). Although most guano-rich caves are in tropical areas (Ferreira 2019; Piló et al. 2023), they also exist in temperate regions, but guano deposition decreases dramatically during winter because of the different seasonal activity of bats (Decu and Tufescu 1976; Gnaspini and Trajano 2000; Ferreira 2019). Studies show that macroinvertebrates often thrive in guano deposits, forming assemblages composed of so-called “guanobiont” species. However, most evidence to this phenomenon comes from tropical regions (e.g., Ferreira and Martins 1999; Ferreira et al. 2000; Moulds 2004; Iskali and Zhang 2015; but see Parenzan and Belmonte 2002; Kováč et al. 2005). Moreover, long-term community-level studies are lacking, and the relationship between macroinvertebrates and environmental variables in guano-rich caves remains poorly understood.

Alpha and beta diversity are essential metrics for studying community structure and biodiversity changes and therefore are routinely used to understand relationship between biodiversity and environmental variables within subterranean ecosystems (Zagmajster et al. 2014; Zagmajster and Christman 2019; Mammola et al. 2024). In its broadest definition, alpha diversity is any measure of biodiversity (e.g., species richness, evenness) within a specific grouping unit (e.g., microhabitats, habitats, communities) at a given scale. In contrast, beta diversity captures species turnover between grouping units or along environmental gradients. Beta diversity can be broken down into specific components such as species replacement (replacement of species between grouping units) and richness differences (loss/gain of species between grouping units) (Cardoso et al. 2014, 2015). At the level of a single cave, a combination of analyses at the alpha and beta levels may allow us to understand what specific assemblages thrive in different microhabitats or specific areas of the cave, and what are the main factors driving biodiversity change across these different areas/microhabitats.

The aim of this work is to characterise the terrestrial macroinvertebrate assemblage inhabiting a temperate guano-rich cave, and to investigate the relationship between environmental variables and structural changes in alpha and beta diversity of the assemblage. We hypothesize bat guano to be the main factor shaping the spatial and temporal distribution of the community, and other abiotic factors to exert a minor effect. Parallel to this primary, ecological-oriented question, we collected novel data on the composition of the community, offering a first checklist of the species of macroinvertebrates inhabiting the investigated cave.

## Methods

### Study area

The cave “Buca dei Ladri” (speleological cataster number: 262 T/Pi) is located in Agnano, Tuscany, Italy. The entrance (43.739139° North, 10.475250° East; WGS84 decimal degree reference system) opens at the foot of the Monti Pisani hills which, in this area, are characterized by the overlap of limestone rocks (dated to Upper Triassic) on metamorphic silico-clastic ones (dated to the transition between Middle - Upper Triassic). The cave develops in Rhetic Limestone, dated to Upper Triassic (Rau and Tongiorgi 1974; Bianucci 1980). The surrounding vegetation includes trees (as *Quercus ilex* L., *Fraxinus ornus* L. subsp. *ornus*), brushes and grass (as *Rhamnus alaternus* L. subsp. *alaternus*, *Phillyrea latifolia* L., *Ruscus aculeatus* L., *Smilax aspera* L.) typical of Mediterranean scrub.

The cave starts with a short horizontal entrance, leading to a 23 meters deep vertical pit. Although not fully appreciable from the map (Figure 1), the cave shows a morphological subdivision into different rooms. At the bottom of the pit, there are two little spaces created by a large rock of collapse present at the base and arranged in an east-west direction; this boulder (referred to as collapsed rock in Figure 1) is just under 10 m tall and therefore the area at the bottom of the pit is unique. After the two rooms, the cave opens up in a larger chamber, which descends through three large terraces to a 11-m deep phreatic lake, located at the level of the aquifer (Bianucci 1980). We considered these five areas in the characterisation of the macroinvertebrate assemblage.

**Fig. 1.**
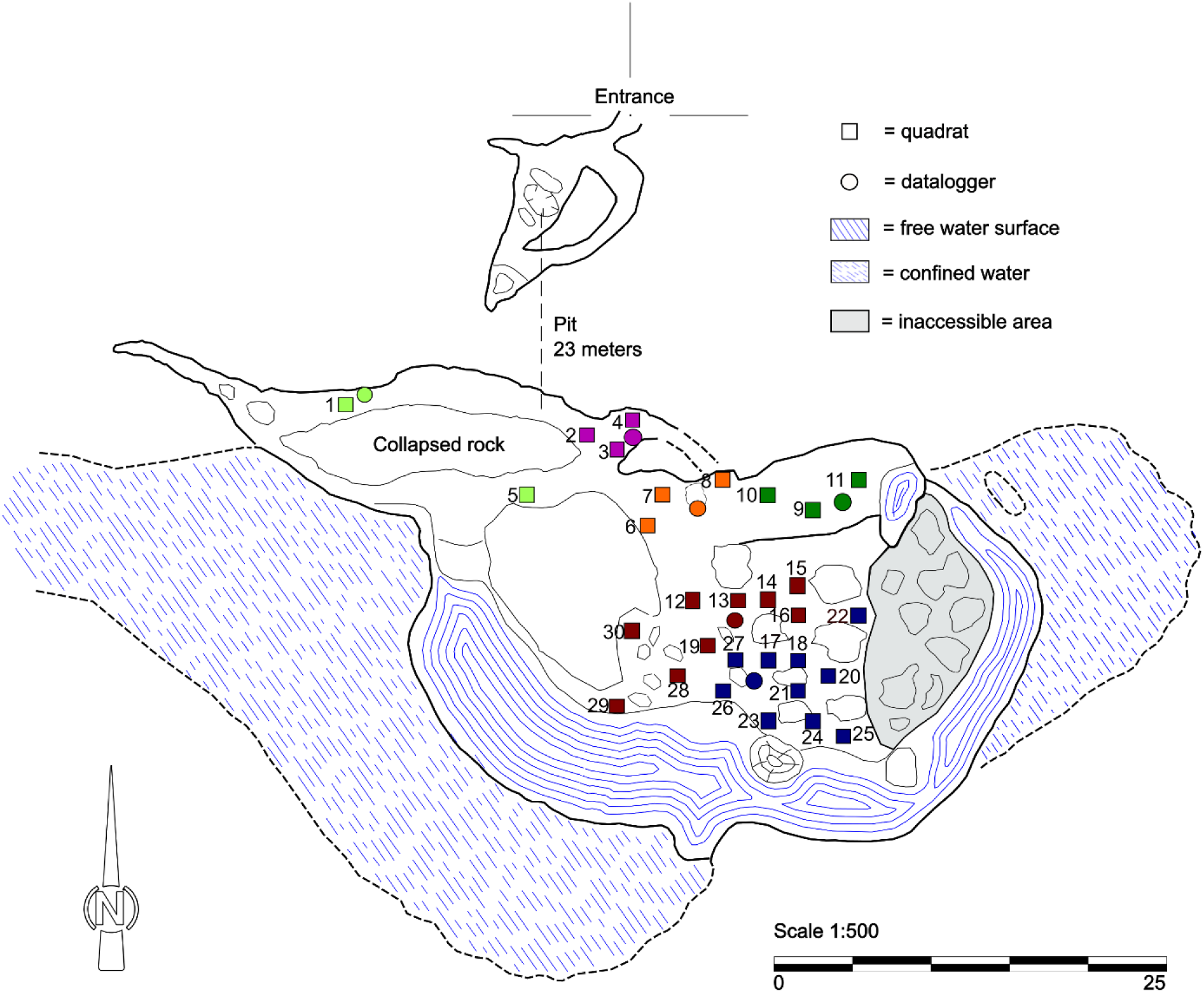
Map of the cave “Buca dei Ladri” with the position of dataloggers and sampling quadrats. The circles indicate the 6 dataloggers (drawn not in scale), the squares the 30 sampling quadrats. The quadrats are drawn in the same colour as the datalogger they were associated with, each colour representing a morphological area of the cave. The grey coloured part represents an inaccessible area due to the instability and slipperiness of the boulders (modified from Federazione Speleologica Toscana 2024. Catasto Grotte Online. Online at: https://www.speleotoscana.it/catasto-online-fst-informazioni/ (accessed on 22/05/24)

The morphology of the entrance, consisting of a short horizontal tunnel leading to the vertical pit, limits the external inputs of organic matter into the cave, mostly restricted to small amounts of vegetable debris at the bottom of the pit. This would make the cave a typical oligotrophic cavity of temperate latitudes; however, during spring and summer, the main room of the cave hosts a large bat colony, including at least three different species (*Myotis blythii* (Tomes, 1857); *Miniopterus schreibersii* Kuhl, 1817; *Rhinolophus ferrumequinum* Schreber, 1774) (P. Agnelli, *pers. comm*. 2023). The seasonal presence of bats caused a huge amount of bat guano to accumulate in the main room of the cave, with the addition of fresh guano in the warmer season.

### Macroinvertebrate identification

Before the beginning of the monitoring and in some cases during the monitoring sessions, we hand-collected 1–5 individuals of each morphospecies for identification. We preserved some of these samples in 70% ethanol and sent them to experts on the different taxonomic groups for morphological identification, whereas we preserved others in 96% ethanol at –20 °C for DNA extraction. Whenever possible, following Balestra et al. (2021) we also took macrophotographs of each morphospecies in its natural environment. In addition, individuals of *Oxychilus meridionalis* from the Apuan Alps, close to the type locality, were included to compare their identity with the individuals from the Buca dei Ladri (Manganelli et al. 2004).

We carried out DNA extraction with the NucleoSpin Plant II DNA kit (Macherey–Nagel GmbH and Co., Düren NRW, Germany) following the manufacturer’s instructions. We conducted the molecular characterisation through sequencing of a 1700–3000 bp fragment of 18S rDNA (henceforth 18S) and Folmer’s fragment (600–700 bp) of cytochrome *c* oxidase subunit I (henceforth COI). We obtained PCR reactions in a 40 μl volume for both markers; a negative control was included in each reaction. We checked amplification success through electrophoresis on agarose gel, all PCR products were purified with the Eurogold Cycle-Pure Kit (EuroClone, Milan, Italy) following the manufacturer’s instructions and sequenced by Azenta Life Service Company. Amplification primers (Medlin et al. 1988; Folmer et al. 1994; Petroni et al. 2002; Lobo et al. 2013) and conditions are listed in Supplementary Material S1, while sequencing primers (Folmer et al. 1994; Lobo et al. 2013; Rosati et al. 2004; Andreoli et al. 2009) are listed in Supplementary Material S2. The sequences obtained have been compared at the level of identity percentage with the ones present on the GenBank and BOLD databases.

### Yearlong monitoring

We monitored the macroinvertebrate assemblage of the cave over one year, using a ‘sampling quadrat’ monitoring approach, as this has been successfully used in previous ecological studies of invertebrate communities in temperate caves (e.g., Mammola and Isaia 2014, 2016, 2018; Mammola et al. 2016a; Balestra et al. 2021). Specifically, we haphazardly placed thirty 1 m^2^ monitoring quadrats on the floors of the cave. We tried to cover the cave surface as homogeneously as possible, while taking into account the presence of rocks and slippery substrates, which makes a standardizing monitoring challenging (Zagmajster et al. 2010; Mammola et al. 2021) (Figure 1). We delimited the corners of each quadrat with four pegs. In each of the morphological areas identified within the cave, we placed a temperature datalogger *Onset HOBO Bluetooth Pendant Temperature* (accuracy: ± 0.5°C), for a total of six dataloggers in the entire cave. We set each datalogger to record a temperature measurement every 30 minutes throughout the study. We georeferenced the position of each quadrat in relation to the entrance of the cave with QGIS version 3.28.4 (QGIS Geographic Information System. Open-Source Geospatial Foundation Project).

We carried out a non-invasive standardized visual census monthly from February 2022 to January 2023. At each visit, a single observer (L.D.) counted the abundance of each morphospecies on the surface of each quadrat for four minutes. The quadrats were not illuminated before the observation to avoid potential interferences with the natural behaviour of the animals. We monitored only the surface of each quadrat, without digging into the substrate or moving any rocks. Hence, we primarily detected morphospecies that spend most of their time on the surface of the substrate, rather than fossorial species.

We expressed the diversity of morphospecies within each quadrat and sampling session (alpha diversity) as the Shannon-Wiener’s diversity index (Shannon and Weaver 1949). This index takes account for the number of species and their abundances. In each sampling quadrat, we also recorded the following independent variables:

1) temperature of the substrate, measured through a thermo-hygrometer TROTEC BC06 (accuracy: ± 0.5°C) The thermometer records the temperature of about 10 cm above the substrate (variable “Substrate temperature”);
2) percentage coverage of the quadrat area in guano (“% Guano”), rock (“% Rock”), soil (“% Soil”), mud (“% Mud”), and water (“% Water”). We measured percentages by splitting each quadrat into 10 cm^2^ sub-quadrats, counting how many sub-quadrats were covered by each substrate (each sub-quadrat roughly representing 1% of coverage). When multiple substrates were represented inside a single sub-quadrat, we considered the most abundant one. The percentages of coverage of the quadrats (% Guano, % Rock, % Soil, % Mud and % Water) were not balanced amongst these categories and were cause of collinearity in the model check. Moreover, during the year it was almost never possible to record percentage changes in the coverage of the quadrats because the guano was often laid in small quantities and in a sparse manner, so the percentage of each substrate remained unchanged in most cases. For these reasons, the percentage coverage was transformed into the categorical variable “Dominant substrate”, obtained by assigning to each quadrat, for each observation, the dominant substrate, defined as covering the quadrat by more than 50%. The levels of this new categorical variable were guano, rock, soil, mud;
3) presence or absence of fresh guano (“Fresh guano”);
4) structural complexity of the quadrat (“Structural complexity”), measured as the length of the diagonal of the quadrat, measured by flattening a soft measuring tape on the diagonal in order to cover the structure of the substrate. This variable represents a proxy of substrate structural complexity (i.e., the longer the linear development, the higher the structural complexity; Mammola et al. 2016a). Note we measured both diagonals, and kept the maximum variable as both measures where highly correlated (Pearson’s *r* = 0.616);
5) depth of the guano (“Guano depth”), as a descriptive indication of the guano already present in the cave. We obtained this measurement inserting a graduated stick in the central part of the guano layer);
6) range of temperature (“Temperature range”) and 7) average temperature (“Average temperature”), obtained retrospectively from datalogger measurements recorded by the datalogger associated with a given quadrat during the monitoring day. Specifically, we associated each quadrat with a specific datalogger based on the quadrat placement inside the morphological areas within the cave (see Figure 1); we verified this spatial association through correlation (Pearson’s |*r|* > 0.6) between measures recorded by the thermo-hygrometer (variable “Substrate temperature” above) and the ones recorded by the associated datalogger.

At the whole cave level, we also marked:

7) bat presence or absence (“Presence of bats”) during the monthly survey;
8) season (“Season”).

### Statistical analyses

We carried out all statistical analysis with the software R version 4.2.2 (R Development Core Team 2022). For graphic elaborations we used the R package “ggplot2” version 3.4.1 (Wickham 2016).

We used a kernel density estimate to visually map the spatial distribution of each species within the cave (Węglarczyk 2018). We estimated both the distribution over the entire year and for monitoring sessions with and without the presence of bats.

To characterise the terrestrial macroinvertebrate assemblage inhabiting the cave, we analysed the role of different explanatory variables in driving changes in alpha and beta diversity in and across sampling quadrats.

At the alpha diversity level, we modelled the association between independent variables and Shannon-Wiener’s diversity index of each quadrat and sampling session through a linear mixed model, fitted via the R package “glmmTMB” version 1.1.5 (Brooks et al. 2017). Prior to model fitting, we explored the dataset following the standard protocol for data exploration by Zuur et al. (2010). In details, we checked the distribution of both categorical and continuous variables, the presence of outliers and multicollinearity among predictors. We evaluated the distribution of continuous variables and the presence of outliers through dot chart. To verify that the levels of the categorical variables were balanced, we constructed bar charts and, if the levels were unbalanced, we merged similar levels of the factors. We investigated multi-collinearity among continuous covariates via Pearson correlation tests, setting the threshold for collinearity at |*r|* > 0.6. We graphically evaluated the association between continuous and categorical variables with boxplots. We dropped collinear variables based on common sense or their biological meaning (Zuur et al. 2010).

We found that five pairs of variables were collinear: Average temperature and Substrate temperature; Season with Substrate temperature, Fresh guano and Bat presence; Presence of bats with Fresh guano (Supplementary Material S3). We decided to keep Substrate temperature, instead of Average temperature, because the former is a single measure for each observation and is not repeated, while the second is the same measure for more than one observation, given that each datalogger is associated with more than one quadrat. Between “Bat presence” and “Fresh guano” we selected the former because it presented more balanced levels. The structure of the resulting model (in R notation) was:

Shannon-Wiener’s diversity index ∼ Substrate temperature + Temperature range + Structural complexity + Presence of bats + Substrate type + (1 | Quadrat) + (1 | Sampling)

Note that we scaled all continuous variables (subtracting the average and dividing by the standard deviation for each value) to facilitate the convergence of the model and ensure the comparability of the effects. The mixed procedure accounted for pseudoreplication due to multiple observations from the same sampling quadrat and multiple observations in same day, by including the spatial variable Quadrat and the temporal variable Sampling as random factors. We validated the model through the function *check_model()* of the R package “performance” version 0.10.2 (Lüdecke et al. 2021), which generates diagnostic plots showing the overlap between observed and predicted values, homogeneity of residuals variance, normality of residuals, normality of random effects, linearity of residuals and collinearity between covariates.

At the beta diversity level, we calculated dissimilarity in species composition between any two quadrats (beta diversity) using the function *beta* of the R package “BAT” version 2.9.2 (Cardoso et al. 2015). Specifically, we expressed community dissimilarity (ranging between zero and one) for pairs of quadrats using the Sørensen index. This function further decomposed total beta diversity into the two underlying components underlying overall dissimilarity (Carvalho et al. 2012): the replacement of species between quadrats (Beta replacement) and the absolute difference between the number of species that each quadrat contains (Beta richness).

We modelled variation in species dissimilarity among quadrats through generalized dissimilarity modelling (Ferrier et al. 2007), as implemented in the R package “gdm” version 1.5.0-9.1 (Mokany et al. 2022). Generalized dissimilarity modelling is a matrix regression technique suitable for modelling variation in beta diversity along gradients. In contrast with standard linear matrix regressions, a generalized dissimilarity model can accommodate for the nonlinear relations between observed compositional dissimilarity and ecological distance between sampling units (for a detailed overview of this approach see Mokany et al. 2022). We constructed three generalized dissimilarity models, relating the three beta diversity matrices (total, replacement and richness difference) to the explanatory variables, allowing us to identify the variables that drive patterns of beta diversity (Mammola et al. 2019a; Mokany et al. 2022). The explanatory variables chosen for the model, based on the data exploration explained above, were: Structural complexity, Spatial distance (distance between quadrats, calculated using the georeferenced map of the cave), Temperature range, Sampling (the date of each sampling session), Substrate temperature, Presence of bats, Rock substrate, Guano substrate, Soil substrate and Mud substrate. The nonlinear patterns were captured using I-splines constructed based on three knots at the default position as defined by the “gdm” package (i.e., minimum, mean and maximum predictor values).

## RESULTS

### Macroinvertebrate identification

We recorded a total of 31 morphospecies of terrestrial macroinvertebrates inside the cave. In detail, we detected 21 morphospecies inside the monitoring quadrats (# in Table 1), 7 morphospecies on the floor outside monitoring quadrats (§ in Table 1), 2 morphospecies on the walls (* in Table 1) and 1 morphospecies parasitizing gastropods of the genus *Oxychilus* (” in Table 1).

**Tab. 1.**
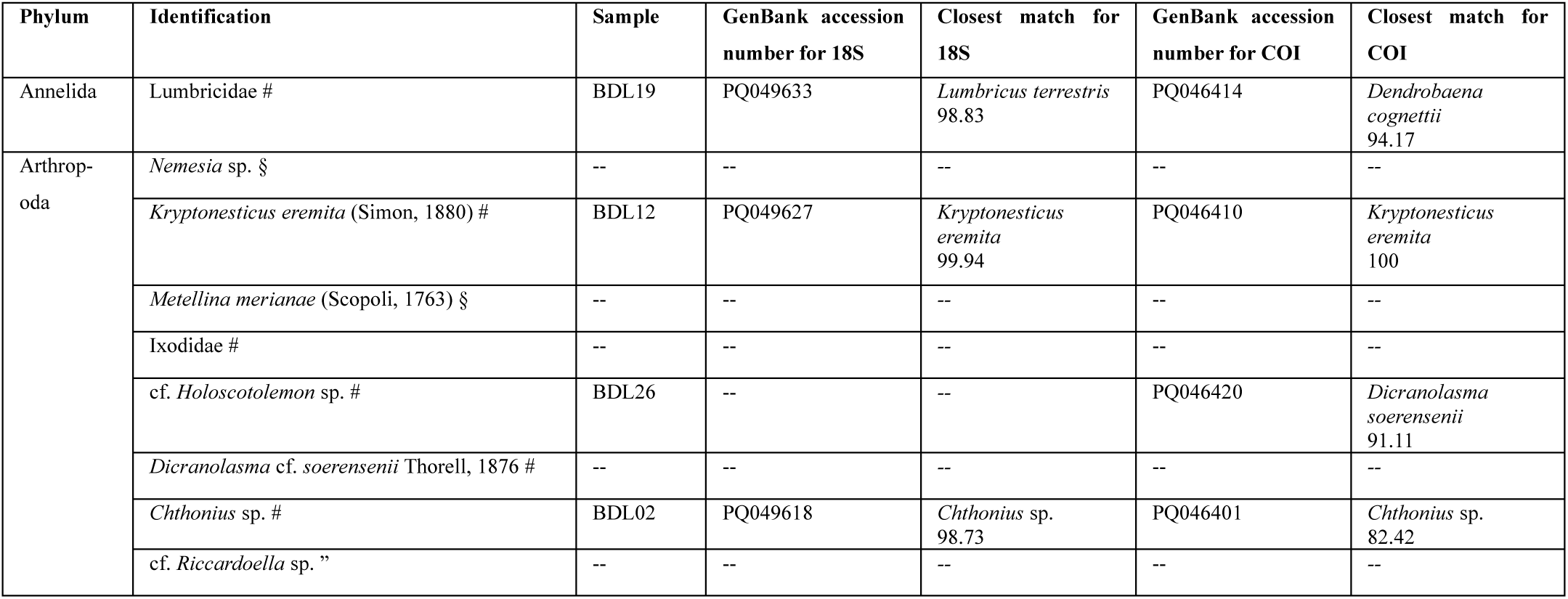
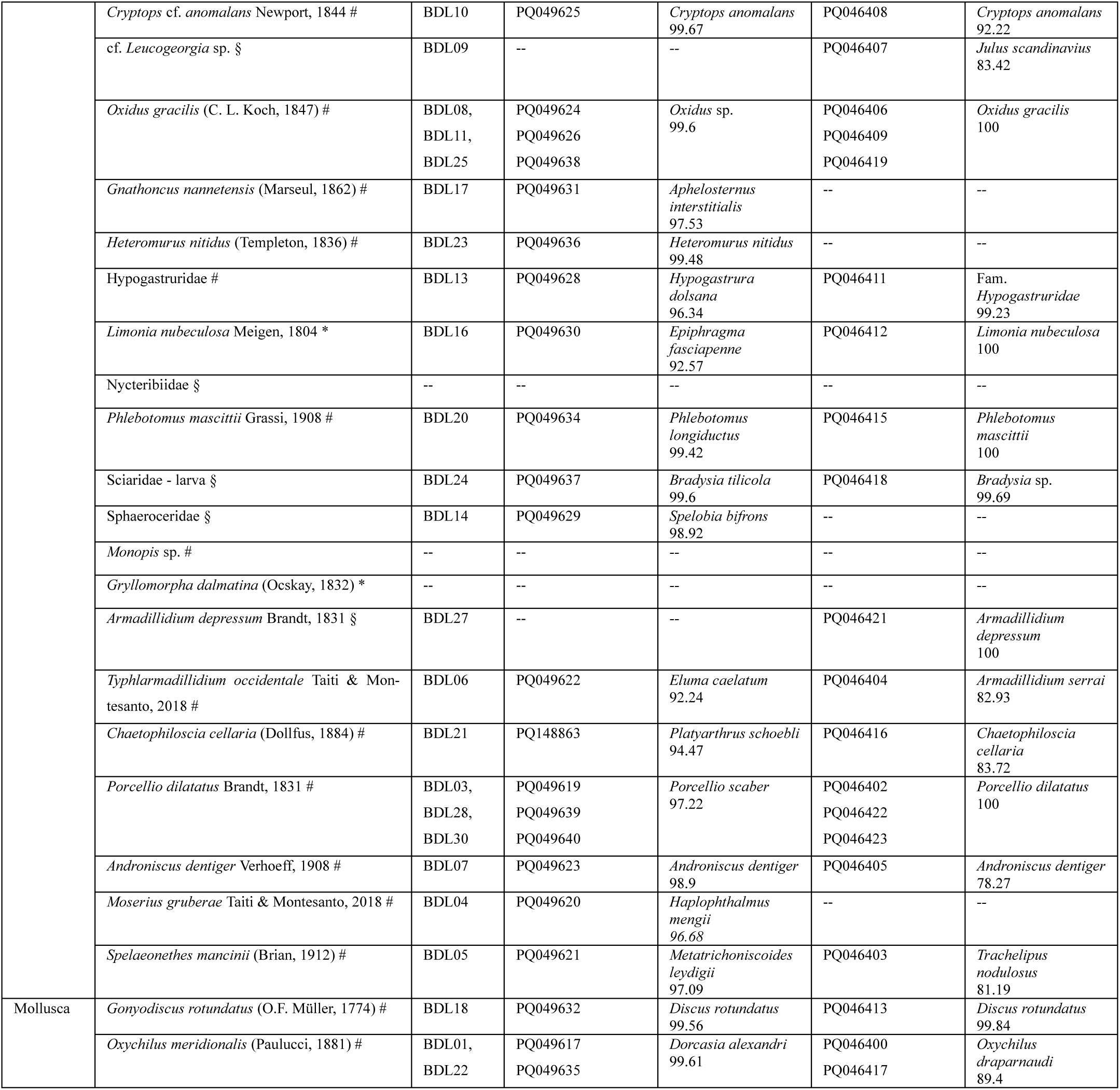
Summary table of the 31 different taxa of terrestrial macroinvertebrates observed inside the cave. The column Identification shows the lowest taxonomic level that could be achieved, the symbol # specifies taxa observed inside the monitoring quadrats, the symbol § specifies taxa observed on the floor outside monitoring quadrats, the symbol * specifies taxa observed on the walls and the symbol ” specifies the taxon observed only in association with gastropods. The column Sample indicates the code used for each sample during molecular characterization. The columns GenBank accession number for 18S/COI give the accession numbers published on GenBank. The column Closest match for 18S specifies the taxonomic information of the sequences with highest percentage of identity with our sequence, obtained from GenBank database. The column Closest match for COI specifies the taxonomic information of the sequences with the highest percentage of identity with our sequence, obtained from GenBank or BOLD database. Nomenclature follows Catalogue of Life (Bánki et al. 2024)

We extracted total genomic DNA from 28 samples, one for each morphospecies. The molecular characterisation allowed us to identify at least 23 MOTUs (Molecular Operational Taxonomic Units), since some specimens initially identified as different morphospecies turned out to be conspecific (see column Sample in Table 1). Moreover, we failed at sequencing for 8 morphospecies (see column Sample in Table 1) The amplification and sequencing of 18S was successful for 25 samples (89%), while for COI was successful for 24 samples (85%). 44% of the 18S sequences obtained showed a >99% percentage of identity with sequences present on GenBank database (Table 1; Figure 2). Regarding COI, 37% of the sequences obtained showed a >97% percentage of identity with the ones present on GenBank; while, using BOLD, 54% of them showed a >97% percentage of identity with the ones present on the database (Table 1; Figure 2).

**Fig. 2.**
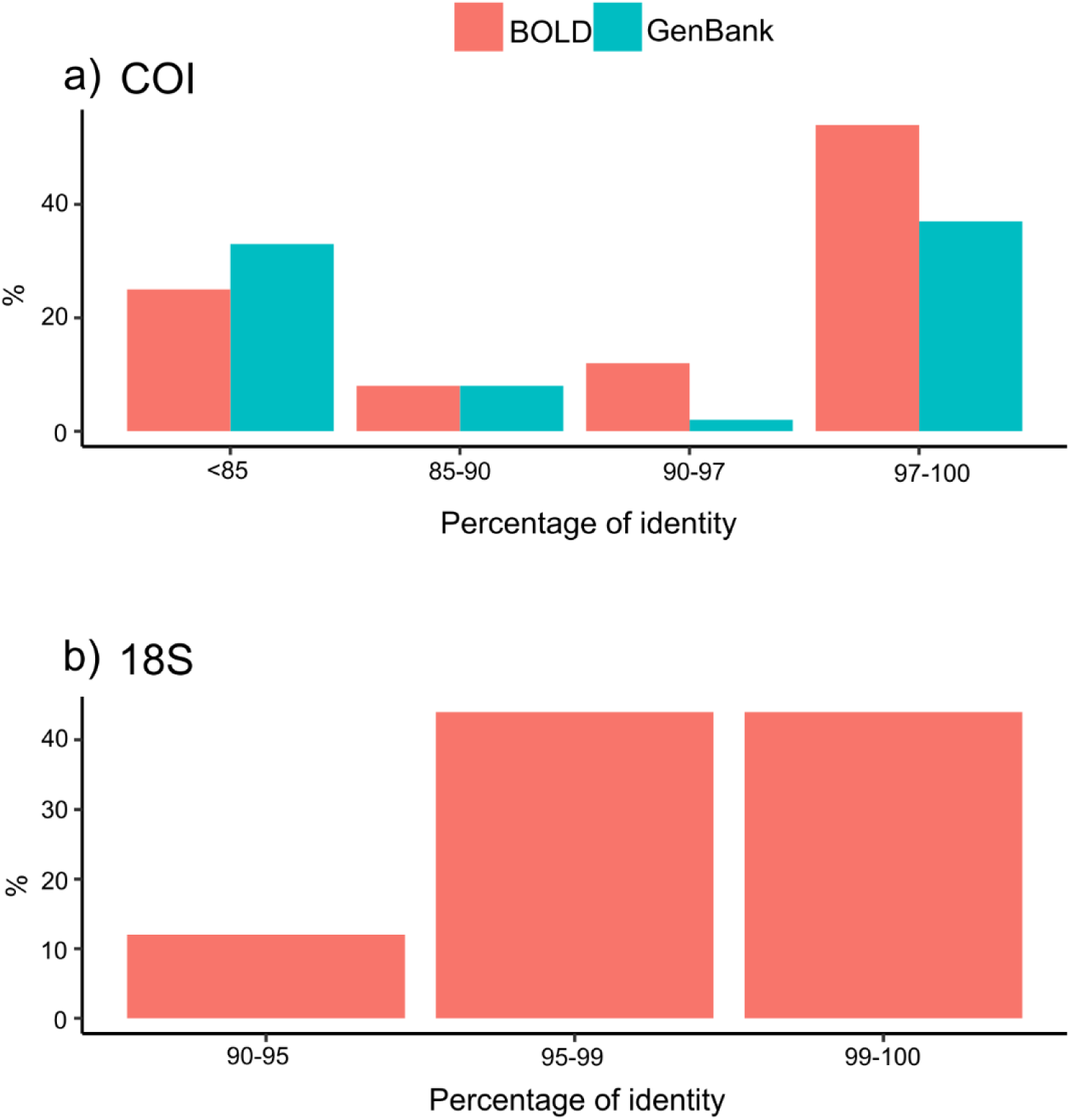
**a**) Percentage of the COI sequences showing a <85%, 85-90%, 90-97% and >97% identity with the ones deposited on GenBank (red) and BOLD (green). **b**) Percentage of 18S sequences showing a 90-95%, 95-99% and <99% identity with sequences present on GenBank

Moreover, we obtained four COI sequences from four individuals of *Oxychilus* cf. *meridionalis* from the Apuan Alps (Supplementary Material S4).

### Cave monitoring

The spatial distributions of species obtained from Kernel estimations showed that the isopod *Porcellio dilatatus* was the only species limited to the first part of the cave (just around the bottom of the pit), and it was present all over the year regardless of the presence or absence of bats (Figure 3, Supplementary Material S5). Collembola *Heteromurus nitidus* (Supplementary Material S6), collembola Hypogastruridae and coleoptera *Gnathoncus nannetensis* had a greater number of observations in the cave when bats were present (Supplementary Material S7).

**Fig. 3.**
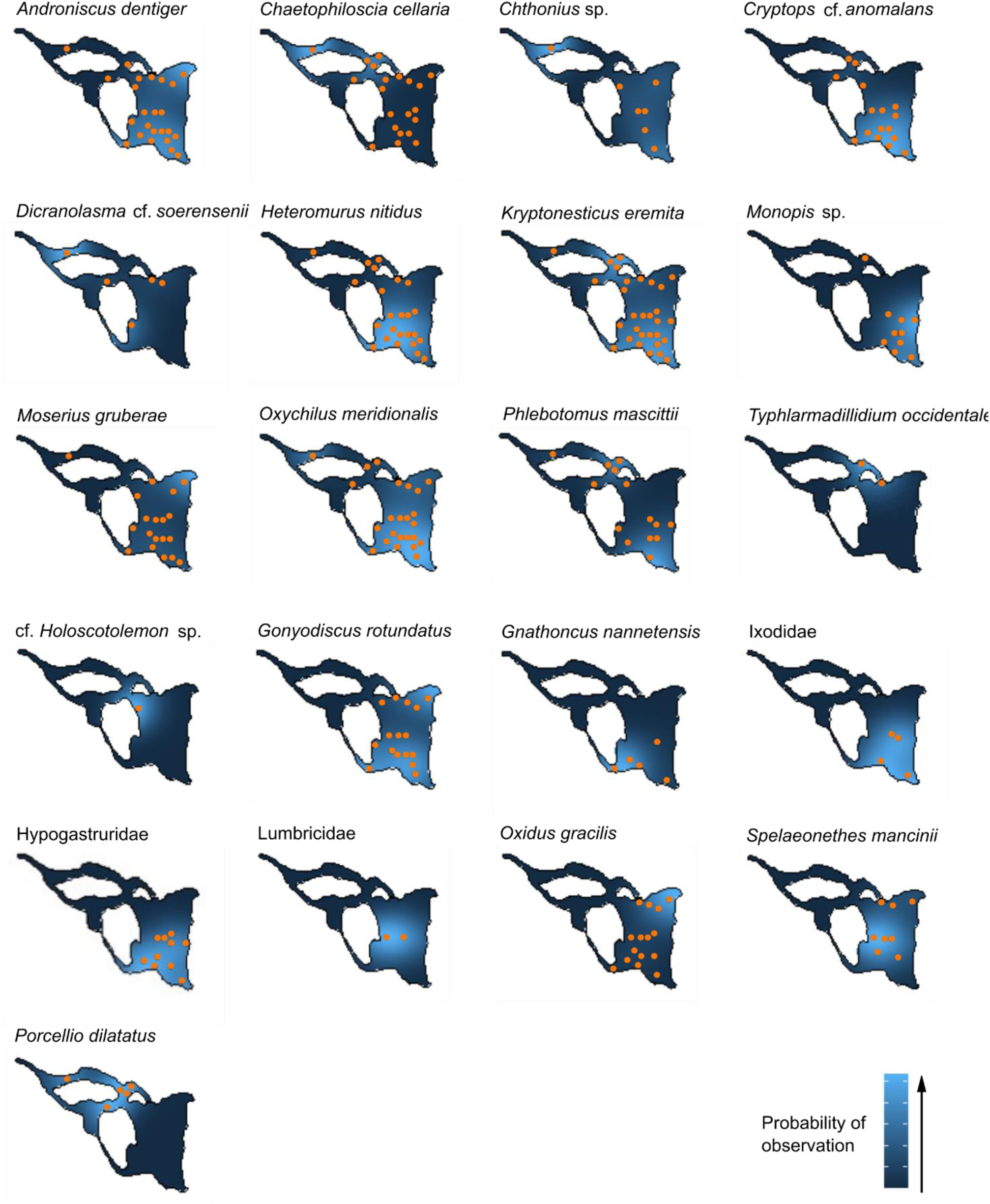
Species distribution inside the cave from Kernel analysis. In each map, the full orange circles indicate the presence of at least one positive observation for that taxon, while the colour scale indicates the probability of observation of a given taxon in a certain portion of the cave. Note that values of the probability of observation are not comparable between different figures, as the maximum and the minimum are calibrated independently for each taxon within the range of variation of the relative abundances

Regarding the descriptive indication of the guano already present in the cave, the depth of the guano varied from 0 to 119 cm in the quadrats considered and showed no noticeable change during the year.

Concerning alpha diversity, species diversity was higher when the bats were present in the cave (estimated β ± standard error [SE] = 0.09 ± 0.04, p = 0.030) and increased with increasing structural complexity (β ± SE = 0.08 ± 0.03, p = 0.015). Species diversity decreased when the dominant substrate was mud (β ± SE = –0.25 ± 0.08, p = 0.036) and soil (β ± SE = –0.21 ± 0.09, p = 0.196) compared to the baseline guano. None of the other variables significantly affected species diversity (Figure 4). The model explained 27% of variance in the data, of which 11% was attributable to the random factors (Figure 4; for model validation see Supplementary Material S8).

**Fig. 4.**
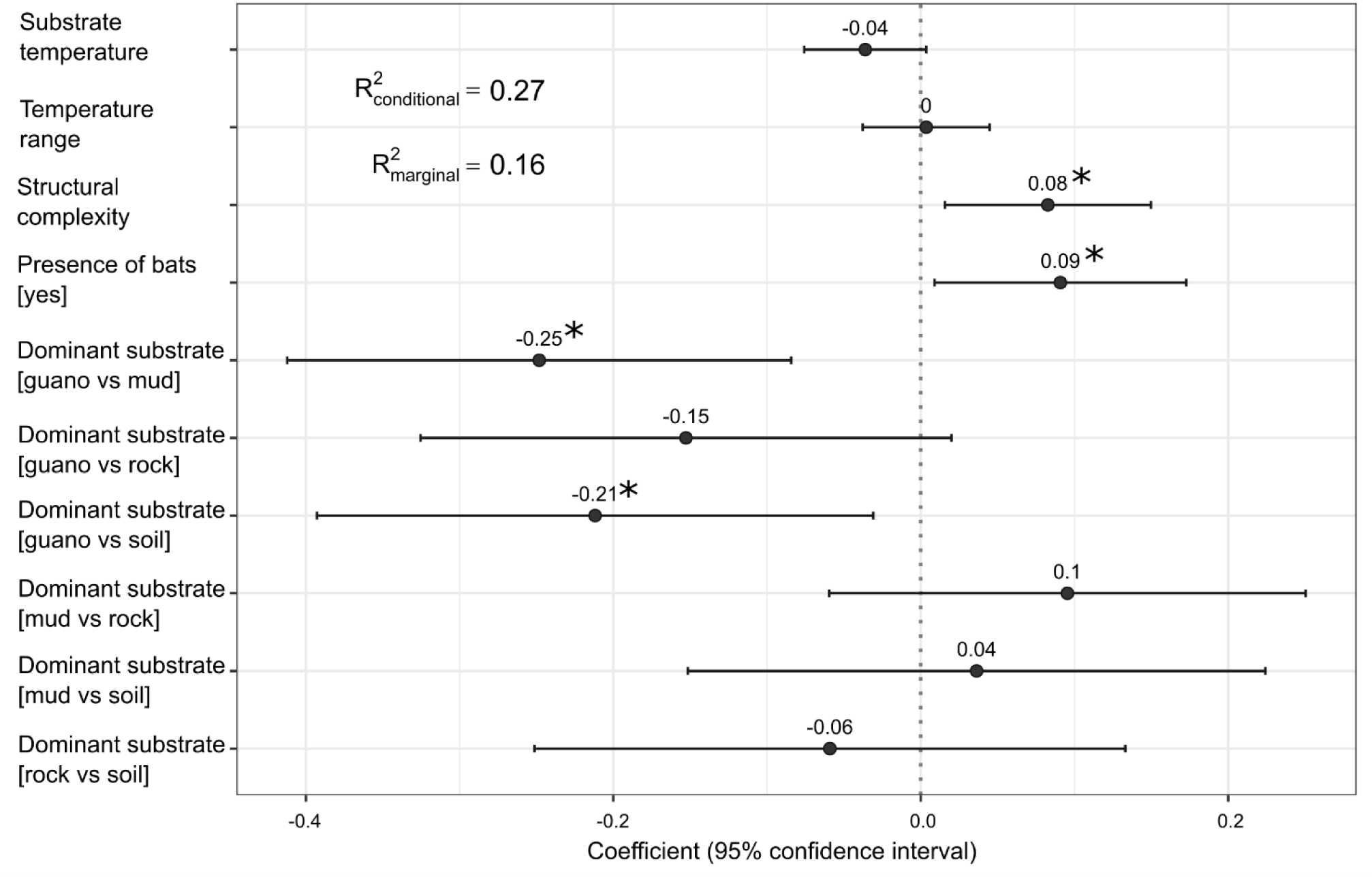
Drivers of species diversity within the cave. Estimated parameters (and 95% confidence interval) for a linear mixed model testing the relationship between species diversity (expressed as the Shannon-Wiener’s diversity index) and five explanatory variables. Asterisks (*) mark significant effects (p < 0.05)

Concerning beta diversity, the structural complexity (Figure 5, panel a) had a greater influence than all other variables in driving species dissimilarity, followed by the spatial distance between quadrats (Figure 5, panel b), substrate temperature (Figure 5, panel e) and temperature range (Figure 5, panel d). The gradient of structural complexity was nonlinearly asymptotic, with rates of turnover steeply increasing between 140 and 150 cm, when it reached a plateau. The gradient of spatial distance between quadrats was nonlinear. The gradient of substrate temperature was exponential and affected the beta diversity starting about from 18 °C. The temperature range affected only the richness different component starting about from 0,5°C of daily thermal excursion The presence of guano (Figure 5, panel h) or mud substrates (Figure 5, panel l) and, to a lesser extent, the presence of bats (Figure 5, panel f), all affected the total beta diversity. Notably, the presence of guano had the strongest effect on total beta diversity and also on the richness difference component, while soil and mud had the strongest effect on the replacement component. However, the most visible effects were that the structural complexity affected the richness difference component while it is negligible regarding the replacement one. Moreover, the distance between quadrats affected the replacement component of beta diversity only when quadrats are further than 20 meters.

**Fig. 5.**
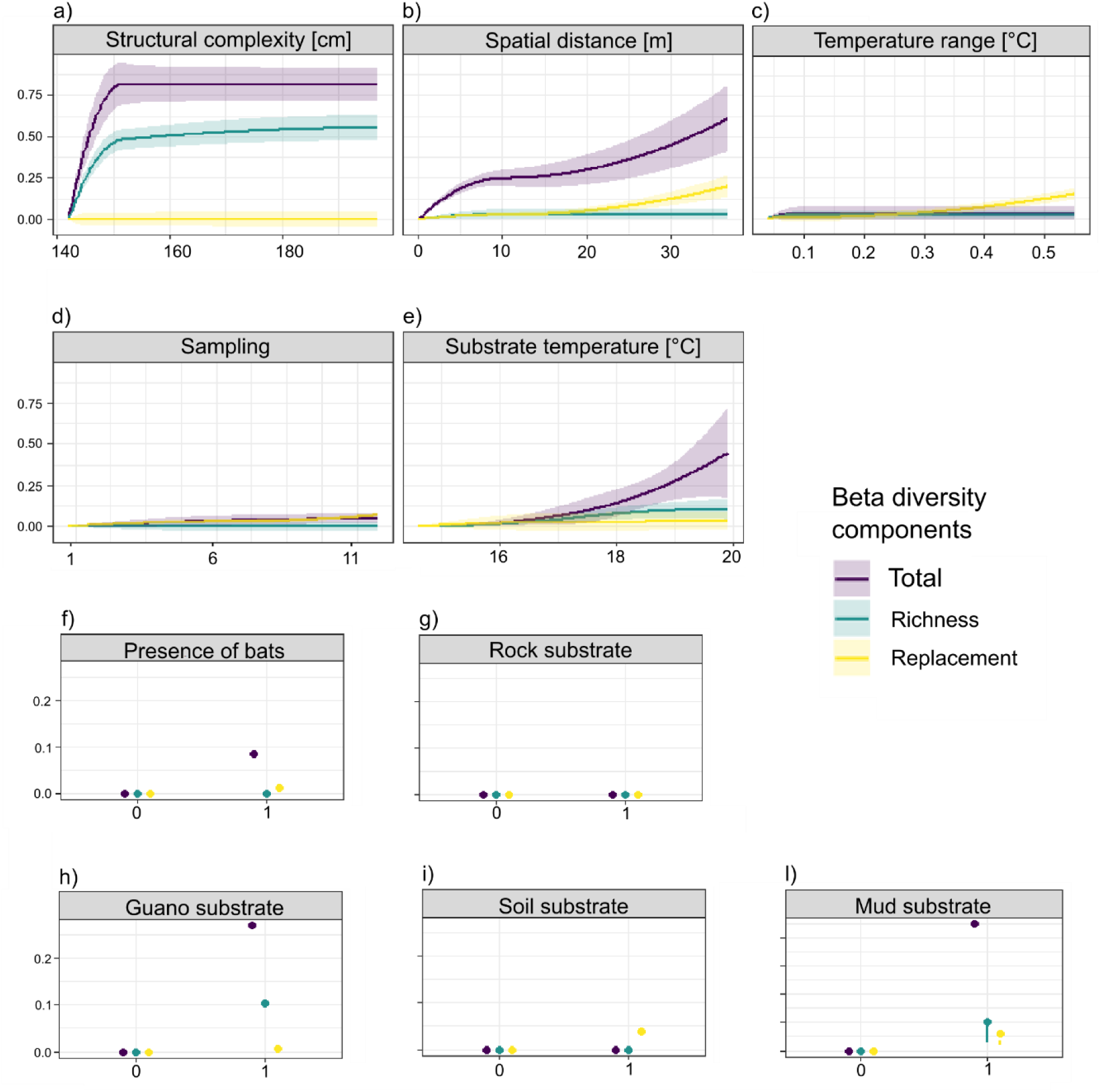
Drivers of species dissimilarity within the cave. On the x-axis is reported the variable with original values, on the y are reported the values transformed by the model. The maximum height reached by each curve provides an indication of the change of β diversity associated with the environmental gradient given by the variable under consideration, holding all other variables of the model constant; the larger the value in y, the more the influence of the variable. In addition, the slope of each function indicates how this influence varies along the gradient of the considered variable (sections of the function with greater inclination imply a greater dissimilarity per unit of the predictive variable), so the shape of each function indicates how the rate of compositional turnover varies along the environmental gradient. As for the categorical variables, the graphic output is a dot, with zero abscissa for the absence and 1 for the presence

## Discussion

### The macroinvertebrate assemblage of the Buca dei Ladri

One year of repeated observations allowed to reconstruct the changes in diversity of the macroinvertebrate assemblage occurring in the Buca dei Ladri, providing its first complete checklist and allowing to identify at least 23 MOTUs belonging to three phyla, which we mostly identified at the genus rank or below. The molecular characterisation of the macroinvertebrates highlighted a limited match with deposited sequences with regard to this fauna: a >97% match on COI sequences, generally considered as indicative of organisms belonging to the same species (Hebert et al. 2003), was obtained on GenBank for 37% of the sequences, on BOLD for 54% of the sequences. Even though the more conserved 18S was considered of limited usefulness for the characterisation of species diversity, as it tends to underestimate it (Tang et al. 2012), also in this case matches with deposited sequences close to 100% represented the 44% of all sequences, highlighting the incomplete status of these libraries. In particular, two specialized isopod species endemic to the Monte Pisano (*Moserius gruberae* and *Typhlarmadillidium occidentale*) and the more widespread *Spelaeonethes mancinii* showed rather low sequence identity (<85% for COI, <97% for 18S) with deposited sequences. This suggests that this group, including several subterranean endemic species with very narrow distributions (Taiti and Montesanto 2018; Taiti et al. 2018; Bedek et al. 2019), lacks enough reference sequences.

Conversely, molecular data highlighted the possible occurrence species complexes in several of the taxa characterised. The most striking case is represented by *Oxychilus meridionalis*, where the sequences from the Buca dei Ladri showed a relatively low identity at the level of the COI (88%) with individuals from the Apuan Alps, close to the type locality, which on their turn belonged to two lineages with a 95% identity (Supplementary Material S9).

A high molecular variability within *O. meridionalis*, suggesting the occurrence of more than one species within this taxon, was already reported by Manganelli et al. (2004) on the basis of ITS1 sequences. The present data support these findings and suggest that *Oxychilus tongiorgii* Giusti, 1969, originally described as an endemic species for the Buca dei Ladri (Giusti 1969) and later synonymised with *O. meridionalis* (Manganelli et al. 2004), might in fact represent a distinct species. The gastropod *Gonyodiscus rotundatus* could also represent a species complex, as our sequence shows 99% identity with sequences deposited in BOLD and 84% identity with sequences deposited in GenBank. A >3% distance with sequences assigned to the same taxon on GenBank and BOLD was retrieved also for *Cryptops anomalans* (92% identity with the closest lineage), *Chaetophiloscia cellaria* (83%) and *Androniscus dentiger* (78%). The last two cases, corresponding to troglophilic isopods with a wide European distribution (Taiti and Ferrara 1995), are particularly interesting, as the identity with allegedly conspecific individuals is extremely low (17-22% distance).

High genetic divergence between allegedly conspecific individuals on mitochondrial markers is well documented in terrestrial isopods (Raupach et al. 2022) and has been explained with several, not fully exclusive hypotheses. Typically, high genetic divergences are interpreted as consequent to phylogeographical events, leading to lineage diversification in allopatry and potentially to speciation (Bedek et al. 2019; Dimitriou et al. 2022; Wang et al. 2024). However, terrestrial isopods are particularly affected by intracellular infection by prokaryotic symbionts of the genus *Wolbachia* (Bouchon et al. 1998), which are known to influence the genetic diversity of mitochondria (Cariou et al. 2017), often leading to discordant patterns between nuclear and mitochondrial genes, and in particular abnormally high intraspecific divergence at the mitochondrial level (Charlat et al. 2009; Jiang et al. 2018). The high genetic diversity identified in *C. cellaria* and *A. dentiger* might therefore be due to these factors, or to less widespread phenomena such as heteroplasmy or atypical mitochondrial DNAs, or to a combination of these factors, and more detailed studies, taking into account a higher number of individuals and populations and including nuclear markers, are needed to ascertain the actual occurrence of a species complex in these taxa.

Interestingly, a non-native species (*Oxidus gracilis*) is one of the most abundant macroinvertebrates in cave (in terms of total abundance it is the fifth species; Supplementary Material S10). *O. gracilis*, with alleged East Asian origin, has currently an almost cosmopolitan distribution; in Europe it was firstly recorded in heavily anthropized environments in the second half of 1800, but is currently widespread in relatively pristine environments (Stoev et al. 2010), including the Monte Pisano area (J. Langeneck, *pers. obs*. 2022). In several parts of its introduced range, *O. gracilis* was reported with reproductive populations in subterranean environments (Esquivel et al. 1986; Reeves 1999; Kunt et al. 2022). Evidence about the impact of non-native species in subterranean ecosystems remain very scarce (Nicolosi et al. 2023; Nicolosi and Gerovasileiou 2024). Our observations, including the record of mating adults and newly hatched juveniles, suggest that *O. gracilis* can complete its life cycle within the cave and is seemingly relevant in terms of biomass, possibly competing with other detritus feeders (the millipede cf. *Leucogeorgia* sp. and several isopods) and being eaten by the larger predators (*Kryptonesticus eremita*, *C. anomalans*).

It’s worth noting that the two cave-dwelling isopods, endemic to the Monte Pisano, *Typhlarmadillidium occidentale* and *Moserius gruberae,* showed quite different abundances. Generally, troglobitic species avoid the larger and fresher guano deposits, while they prefer old guano mixed with soil (Ferreira 2019). Probably, the competition with abundant populations of guanobiont organisms, combined their preference for oligotrophic environments, means that these are usually rare in the presence of guano (Howarth 2023). *T. occidentale* was present with few individuals (2 observations in all monitoring), as is typical of the forms adapted to the cave; *M. gruberae* was instead ubiquitous (250 observations in all monitoring) (Supplementary Material S10). Compared to all other isopod species recorded during the survey, it shows smaller dimensions and a morphology of the exoskeleton rich in tubercles, making it more resistant. The small size of *M. gruberae* could allow it to avoid competition with larger isopods and, together with the particular morphology of the exoskeleton, make it less palatable to predators.

The macroinvertebrate fauna within the Buca dei Ladri mostly includes troglophilic species with a wide distribution encompassing a significant part of Europe (e.g., *K. eremita*, a very common European cave-dwelling spiders; Mammola et al., 2022), with a minor part of allegedly endemic species with a much narrower distribution. However, at least some of the species with an allegedly wide distribution showed the presence of multiple lineages, suggesting that they might represent species complexes, and that their systematics should be reviewed. More generally, while subterranean fauna in Italy has been the subject of several inventories and taxonomic studies (Bertolani et al. 1994; Parenzan and Belmonte, 2002; Isaia and Pantini, 2010; Sabella et al. 2020; Latella 2022), their actual distribution and degree of endemicity is often unclear, and molecular data are particularly scanty and restricted to a few taxa (e.g., Allegrucci et al. 2014; Mammola et al. 2016b; Isaia et al. 2017; Taiti et al. 2018). The comparison of our data with the available repositories highlights a very low coverage of the taxa retrieved in this study, and of the most specialized ones, stressing the need for further molecular studies both taking into consideration the whole macroinvertebrate assemblage occurring in a cave system, and focusing on specific taxa.

### The ecology of a guano-rich temperate cave

We hypothesized bat guano to be the main factor shaping the spatio-temporal distribution of the community, and other abiotic factors to exert a minor effect. Yet, the availability of guano turned out not to be the primary factor driving the community dynamics within the cave. Structural complexity was the main factor positively influencing species diversity, matching previous study on cave-dwelling spiders based on a similar sampling methodology (Mammola et al. 2016a). Our results show that both alpha and beta diversity were largely affected by this predictor, indicating that the observed increase in alpha diversity was mainly driven by the increased colonisation in highly complex quadrats, with no apparent replacement of species. While in most of cases the replacement component leads changes in beta diversity (Graco-Roza et al. 2022), our findings present a different perspective of the processes driving community composition in subterranean environments. Generally, quadrats with a greater structural complexity contain stones, debris and holes, leading to an effective surface area of more than 1 m². We speculate that this availability of spaces provides an increase availability of niches to be exploited by several species, acting as shelters or, in the case of spiders, as anchor points to build 3-dimensional webs (Mammola and Isaia 2017).

In addition, the presence of bats also led to an increase in species diversity. Caves colonised by bats often present fresh guano deposits and ectoparasites (in this particular instance ticks and Nycteribiids) that can enter the trophic network (L.D. *pers. obs*. 2024). While the alpha diversity in quadrats with guano substrate versus ones with mud or soil substrate was smaller, contradicting our working hypothesis, we explain this by the fact that the majority of the guano in the quadrats was old and, as such, scarcely nutritious (Ferreira 2019). In fact, the guano representing the dominant substrate in the first observation (February 2022) was old, as bats had moved out of the cave months before, and during the year none of the quadrats shifted to having guano as the dominant substrate. Fresh guano was detected during the study, but it was always in small patches, scattered on the substrate, and never abundant enough to cover all the surface. Based on our observation, in the sampling year the bat colony was not above the sampling quadrats and seemed to have moved from above the areas with stacks of old guano to the right terminal part of the cave (i.e., an inaccessible part of the cave; Figure 1) and above the lake.

While the strongest effect was observed due to on richness differences in our study, species replacement occurred in quadrats at least 25m apart, in wider temperature ranges and in soil substrates. It is noteworthy that quadrats were only 25m apart when we combined a quadrat from the cave entrance with one placed in the inner part of the cave (Figure 1). Our finding thus emerges from the fact that some species were only found in both ends, for example, *P. dilatatus* was found only in the first part of the cave, while *G. nannetensis* only in the inner part. This result is largely expected, as distance from the surface is a key factor explaining change in species composition within caves (e.g., Gers 1998; Ferreira et al. 2007; Novak et al. 2012; Tobin et al. 2013; Mammola et al. 2017; Kozel et al. 2019). Still, we did not find an effect of spatial distance in the richness differences, probably because Buca dei Ladri presents guano deposits in the innermost parts of the cave and the organic substance load does not decrease according to the external-internal gradient. Interestingly, the majority of the species occurring in the inner part are not exclusive to the subterranean environment and only three isopod species (*Typhlarmadillidium occidentale*, *Moserius gruberae* and *Spelaeonethes mancinii*) showed obvious adaptation to a subterranean lifestyle. A daily thermal excursion of 0.5°C or more also influences the richness component. These values are recorded only in the areas covered by loggers 1 and 2 (Figure 1; Supplementary Material S11), these are the two parts of the cave that, for the circulation of air, lead from entrances (see below for more details). So this data complements and confirms the fact that in the cave there is an effect mainly on the replacement component given by proximity or distance from the outside.

Temperature affected the beta diversity starting from 18 °C (Figure 8E); however, temperatures higher than 18 °C occur only in summer in the cave area covered by datalogger 2 (Figure 1; Supplementary Material S11). This area probably undergoes an external influence at temperature level; from the analysis of the annual trend of temperatures taken by the dataloggers (Supplementary Material S11), it seems that there is a second entrance for the air, probably consisting of cracks in the ceiling, not accessible to humans. Moreover, the guano substrate (Figure 5L) and the mud one (Figure 5H) affected beta diversity (Figure 5F), suggesting distinct contingents of species with preference for different substrate conditions.

Finally, there were variations in species composition driven by the seasonal presence of bats in the cave, confirming that the presence of the bats has a detectable, although small influence on the overall macroinvertebrate assemblage.

### Methodological considerations

Unlike the quadrats defined on the floor of the cave, both the substrate outside the sampling quadrats and the walls of the cave were not sampled in a standardised manner. For this reason, the numbers of respectively seven and two morphospecies found in these areas were probably underestimation and could not be compared with the number of 21 morphospecies detected inside the quadrats.

Setting standardized experimental design in caves is a challenge due to the known unfavourable working conditions (Mammola et al. 2021). The experimental design we used in this study allowed us to collect information and sample most of the cave surface. However, the inaccessible area and the phreatic lake remained out of sight. Additionally, by observing only the surface of the substrate, we were able to make observations comparable to each other and not distorted by further research methods (such as lifting stones, digging moving the soil, or using different trapping). Hence, likely this approach underestimated total biodiversity. Such underestimation already occurs because the cave environment is connected a network of cracks and fissures that animals use, but which we cannot explore. When humans enter a cave, they have an anthropocentric perspective; we enter to study animals, but we can only do so in the spaces accessible to us, which are much fewer than those available to animals (Mammola 2019; Mammola et al. 2021). Nonetheless, a standardized method of monitoring, applied with an understanding of these objective limitations, can help make studies comparable and replicable in different caves.

Studying caves with a standardised methodology is also crucial in a context of climate change. Since these environments are very stable, the recovery time after a disturbance can be expected to be rather long, and the species occurring in these environments typically show low tolerance to climatic disturbances (Mammola et al. 2019b; Vaccarelli et al. 2023). This is particularly true for endemic species which, while not representing the majority of the taxa identified, occur in the Buca dei Ladri. In our case, the fact that the assemblage is related to the presence of bat guano, makes it particularly vulnerable to climate change; indeed, bats make a selection based on the microclimate within their resting sites to minimize the adverse effects of climate change, so their distribution is likely to be changed in the next years (Vaccarelli et al. 2023), with all the cascade effects that result if they were to leave the cave. Unpublished observations (P. Agnelli, *pers. comm.* 2023) already highlighted a noticeable trend of decrease of bat populations in the cave, and this phenomenon might have cascading effects on the remaining biota. In this context, unravelling the relationship between macroinvertebrate assemblages and abiotic variables, particularly temperature, is pivotal to address the effects of the current biodiversity crisis in these sensitive environments, to forecast future scenarios, and to plan conservation measures.

## Conclusion

We provided the first checklist of the macroinvertebrates in Buca dei Ladri cave and first insights into the ecology of guano-rich temperate caves. We found that macroinvertebrate distribution at the alpha and beta diversity levels was mostly explained by the structure of the substrate. Alpha diversity was positively associated with a more complex substrate, regarding beta diversity only richness difference was influenced, unlike what happens in most temperate caves, possibly due to the higher abundance of food sources through the whole extension of the cave. Moreover, bat presence was also associated with an increase in both alpha and beta diversity.

The macroinvertebrate assemblage was primarily characterised by troglophilic, widespread species. We confirm the presence of two Monte Pisano’s endemic isopods highly specialized to a subterranean lifestyle: *Moserius gruberae* and *Typhlarmadillidium occidentale*. Importantly, molecular data suggest that some of the most widespread species are in fact species complexes and would warrant further taxonomic investigations. Conversely, the non-indigenous diplopod *Oxidus gracilis*, originating in eastern Asia, was one of the most abundant species, and the observations carried out suggest that it is able to complete its life cycle within the cave.

Notwithstanding some limitations of the sampling approach, leading to a probable underestimation of the overall cave’s biodiversity, we suggest that integrating a zoological approach, aimed at giving a precise identification of the species, and an ecological approach, focusing on patterns of biodiversity change, is key to better understand the biotic structure of subterranean ecosystems and how they can be affected by future environmental changes.

## Acknowledgements

Research on caves requires the technical support of many speleologists but, at the same time, the involvement of people also spreads a culture of knowledge and respect for the environment, thus supporting biodiversity conservation. We are grateful to all members of the Gruppo Speleologico Archeologico Livornese, particularly to Adriano Civitillo, Andrea Massagli, Claudia Ferro, Daniele Pagli, Davide Viola, Dulia Melluso, Elena Casolaro, Enrico Lauretti, Giulia Bolognini, Giulio Della Croce, Marco Della Mea, Michele Brondi, and Roberta Mezzena. Special thanks go to Giuseppe Mazza, Luigi Romani, Paolo Agnelli, Folco Giusti, Fabio Macchioni, Fabio Penati, Stefano Taiti, and Marzio Zapparoli for their invaluable support in taxonomy identification. We thank Irene Tatini and Sara Verni for their helpfulness in the laboratory.

## Author Contributions

LD, SM, AM, JL, FS, and GP conceived the study. LD, FS, AM, and SM carried out a preliminary inspection of the site. LD, SM, and JL designed methodology. GP provided financial resources. LD conducted fieldwork and curated data. LD, JL and LG carried out molecular analyses. LD, SM and CG-R analysed data. LD wrote the first draft, with substantial contributions by SM, JL, AM, and CG-R. All authors reviewed the manuscript and provided critical suggestions and additions to the text.

## Data Availability

Sequences are available on GenBank (after paper acceptance).

The complete dataset supporting this study is available on Figshare (doi will be provided upon acceptance).

