## Supplementary Materials for "Macroinvertebrate diversity patterns in a guano-rich temperate cave"

S1

S2

S3

S4

S5

S6

S7

S8

S9

S10

S11

### Supplementary materials

**S1.** PCR protocols and primers for each of the amplified markers. For the primer pair designed by Lobo et al. (2013), several attempts were made varying the temperature of annealing (40 ºC, 45 ºC, and 50 ºC)

| Amplified marker | PCR protocol | PCR primer | Primer sequence 5'-3' | Bibliography |
|---|---|---|---|---|
| COI | 3 min 94 °C, 35 × [30 s 95 °C, 60 s 40-50 °C, 60 s 72 °C], 6 min 72 °C. | LCO1490<br>HCO2198<br>LOBO F1<br>LOBO R1 | -GTCAACAAATCATAAAGATATTGG-<br>-TAAACTTCAGGGTGACCAAAAAATCA-<br>-KBTCHACAAAYCAYAARGAYATHGG-<br>-TGRTTYTTYGGWCAYCCWGARGTTTA- | Folmer et al. 1994<br>Folmer et al. 1994<br>Lobo et al. 2013<br>Lobo et al. 2013 |
| 18S | 3 min 94 °C, 35 × [30 s 94 °C, 30 s 54 °C, 120 s 72 °C], 6 min 72 °C. | F9<br>R1513 Hypo | -CTGGTTGATCCTGCGAG-<br>-TGATCCTTCYGCAGGTTC- | Medlin et al. 1988<br>Petroni et al. 2002 |

**S2**. Sequencing primers employed for each of the sequenced markers

| Sequenced marker | Sequencing primer | Primer sequence 5'-3' | Bibliography |
|---|---|---|---|
| COI | HCO1490<br>LCO2198<br>LOBO R1<br>LOBO F1 | -GTCAACAAATCATAAAGATATTGG-<br>-TAAACTTCAGGGTGACCAAAAAATCA-<br>-KBTCHACAAAYCAYAARGAYATHGG-<br>-TGRTTYTTYGGWCAYCCWGARGTTTA- | Folmer et al. 1994<br>Folmer et al. 1994<br>Lobo et al. 2013<br>Lobo et al. 2013 |
| 18S | R536<br>F300<br>R1052<br>F783<br>F919 | -CTGGAATTACCGCGGCTG-<br>-AGGGTTCGATTCCGGAGA-<br>-AACTAAGAACGGCCATGCA-<br>-GACGATCAGATACCGTC-<br>-ATTGACGGAAGGGCACCA- | Rosati et al. 2004<br>Andreoli et al. 2009<br>Rosati et al. 2004<br>Rosati et al. 2004<br>Rosati et al. 2004 |

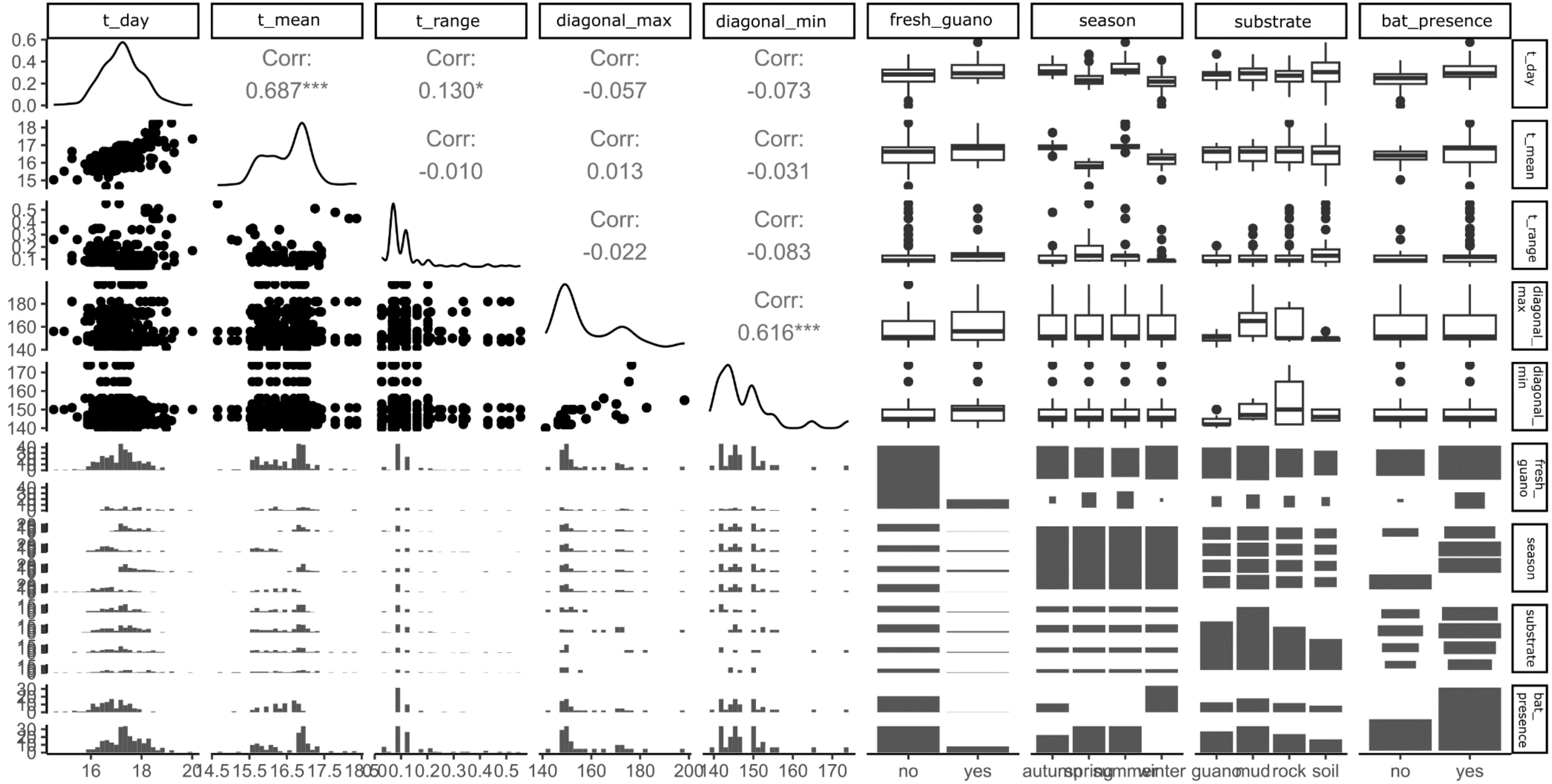


**S3**. Output of the Pearson correlation tests to investigated multi-collinearity among continuous covariates

**S4.** The four sequences obtained from individuals of *O.* cf. *meridionalis* from caves in the Apuan Alps, with relative accession numbers and coordinates, are reported. Moreover, the same data are shown also for *Chilostoma cingulatum apuanum*, used as outgroup in the phylogeny

| **Taxon** | **GenBank accession number** | **Coordinates** |
|---|---|---|
| *O.* cf. *meridionalis* | PQ046427 | 44.107500° N, 10.150556° E |
| *O.* cf. *meridionalis* | PQ046426 | 44.107528° N, 10.151056° E |
| *O.* cf. *meridionalis* | PQ046425 | 43.984211° N, 10.337926° E |
| *O.* cf. *meridionalis* | PQ046424 | 43.971257° N, 10.533177° E |
| *Chilostoma cingulatum apuanum* | PQ046428 | 44.107791° N, 10.150190° E |

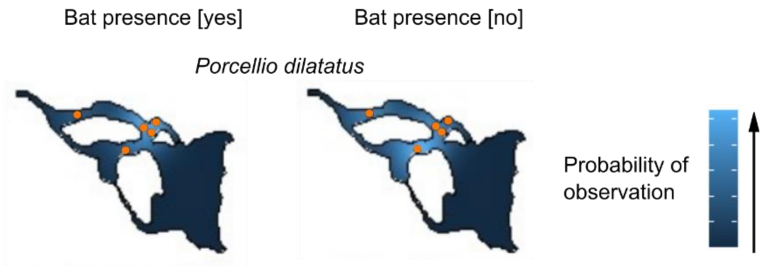


**S5**. *Porcellio dilatatus* distribution inside the cave from Kernel analysis, with associated the data of presence/ absence of bats. It's the only species recorded only in the area at the bottom of the pit. In each map, the full orange circles indicate the presence of at least one positive observation for that taxon, while the colour scale indicates the probability of observation of *P. dilatatus* in a certain portion of the cave

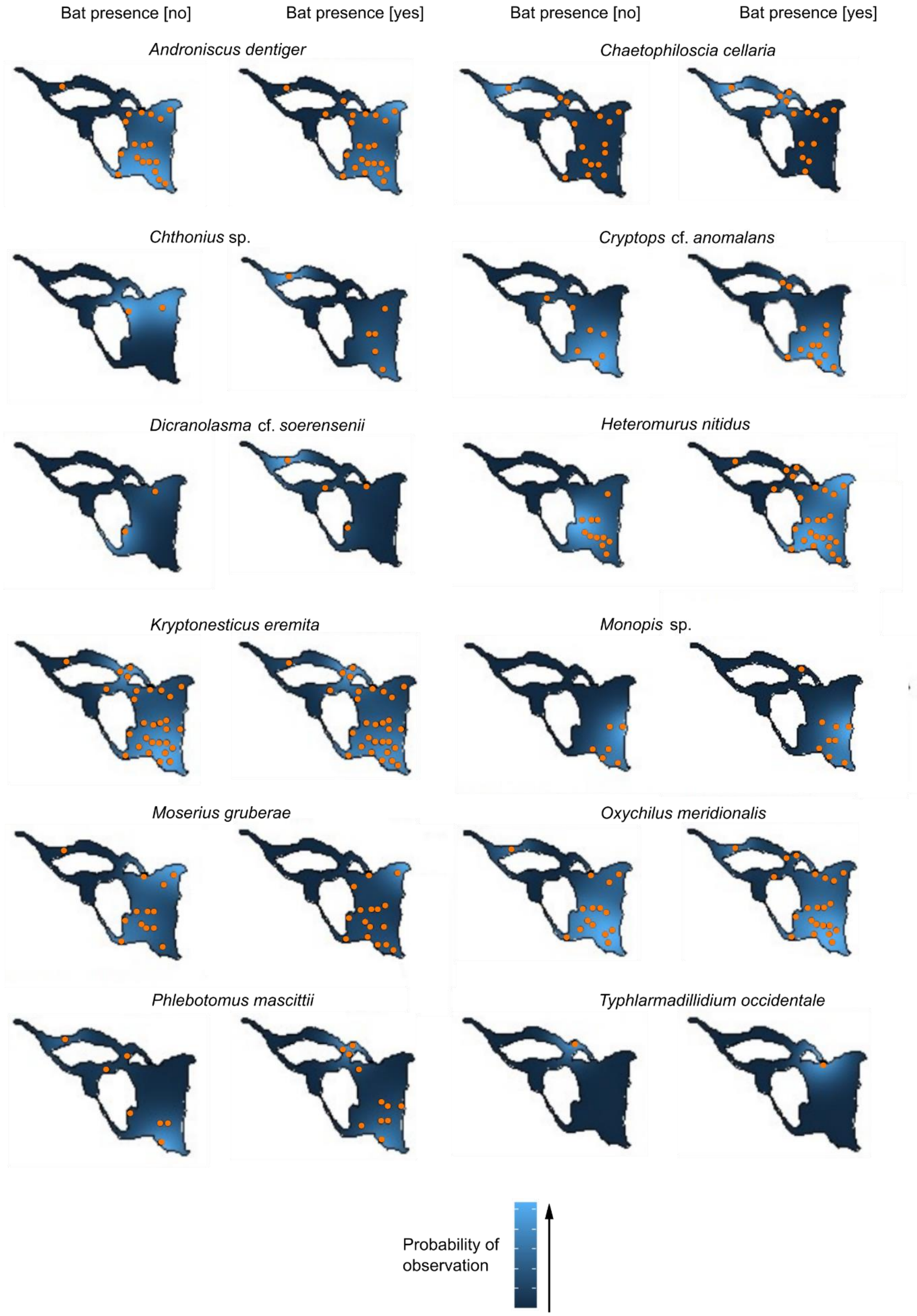


**S6.** Species distribution inside the cave from Kernel analysis, with associated the data of presence/ absence of bats. In this case are reported only taxa present in all the extension of the cave (both at the bottom of the pit and in the rest of the cave). In each map, the full orange circles indicate the presence of at least one positive observation for that taxon, while the colour scale indicates the probability of observation of a given taxon in a certain portion of the cave. Note that values of the probability of observation are not comparable between different figures, as the maximum and the minimum are calibrated independently for each taxon within the range of variation of the relative abundances

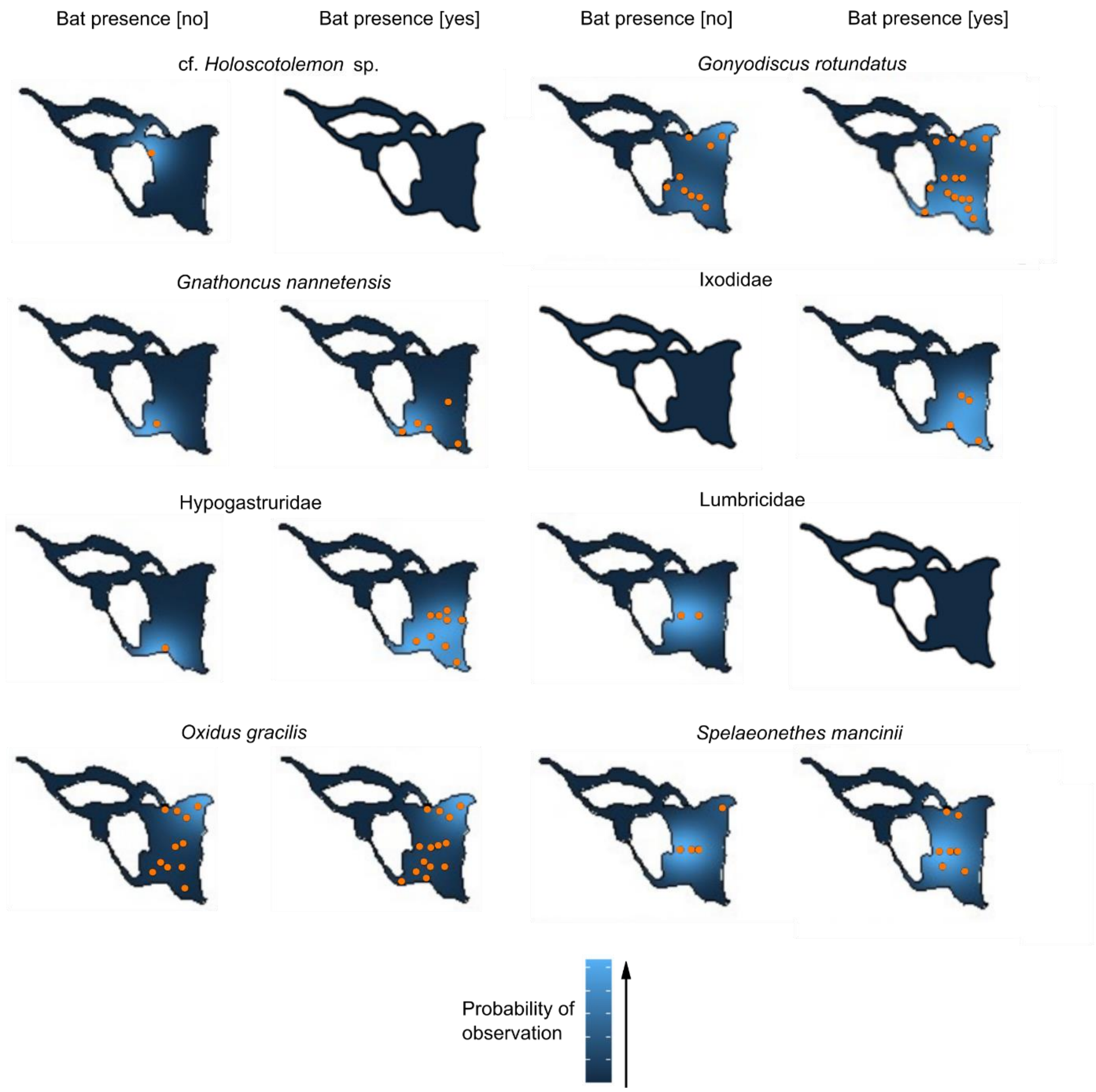


**S7.** Species distribution inside the cave from Kernel analysis, with associated the data of presence/ absence of bats. In this case are reported only taxa not recorded at the bottom of the pit. In each map, the full orange circles indicate the presence of at least one positive observation for that taxon, while the colour scale indicates the probability of observation of a given taxon in a certain portion of the cave. Note that values of the probability of observation are not comparable between different figures, as the maximum and the minimum are calibrated independently for each taxon within the range of variation of the relative abundances

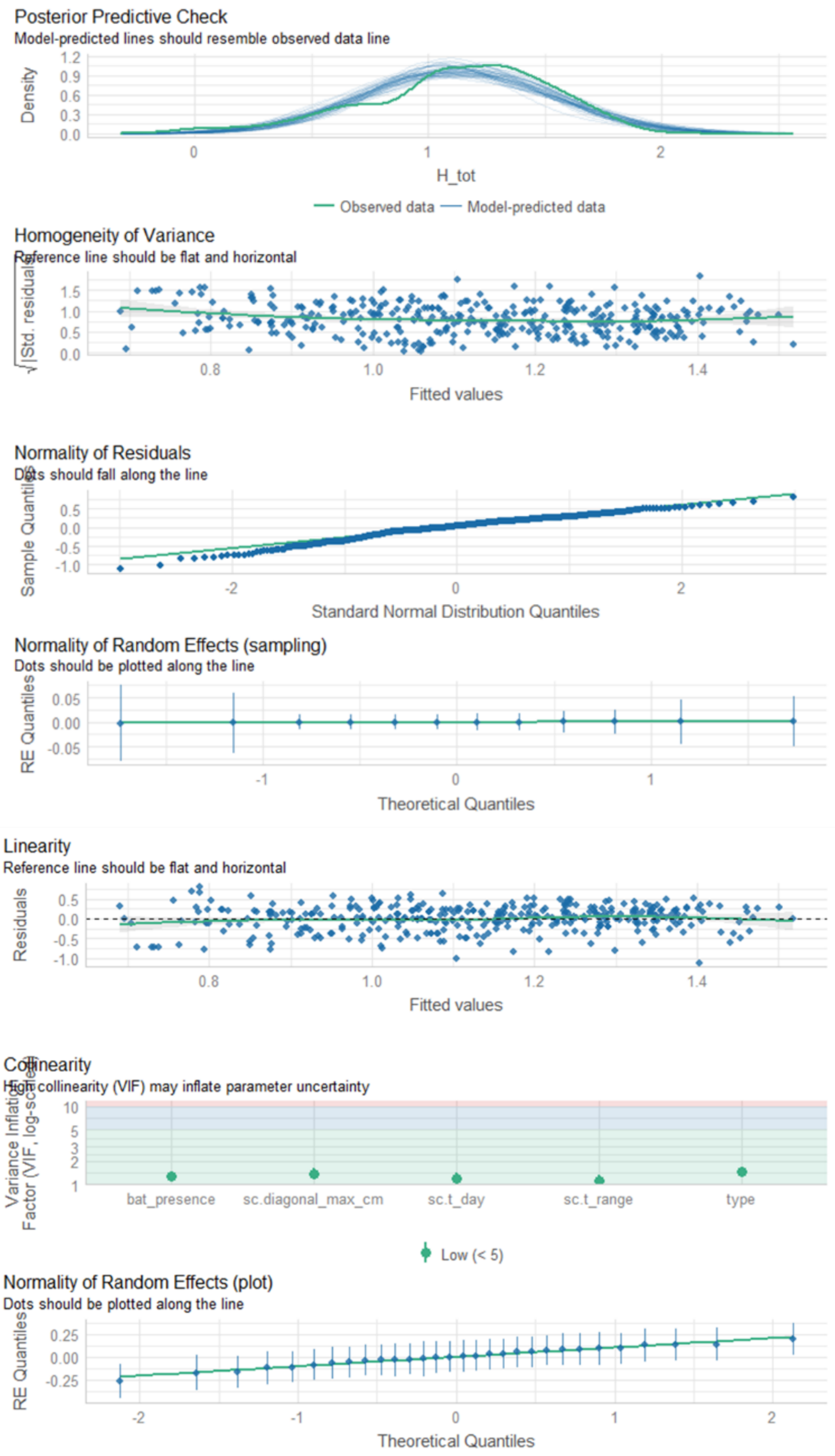


**S8.** Graphic validation of the model using the function *check_model()* of the R package "performance" version 0.10.2 (Lüdecke et al. 2021)

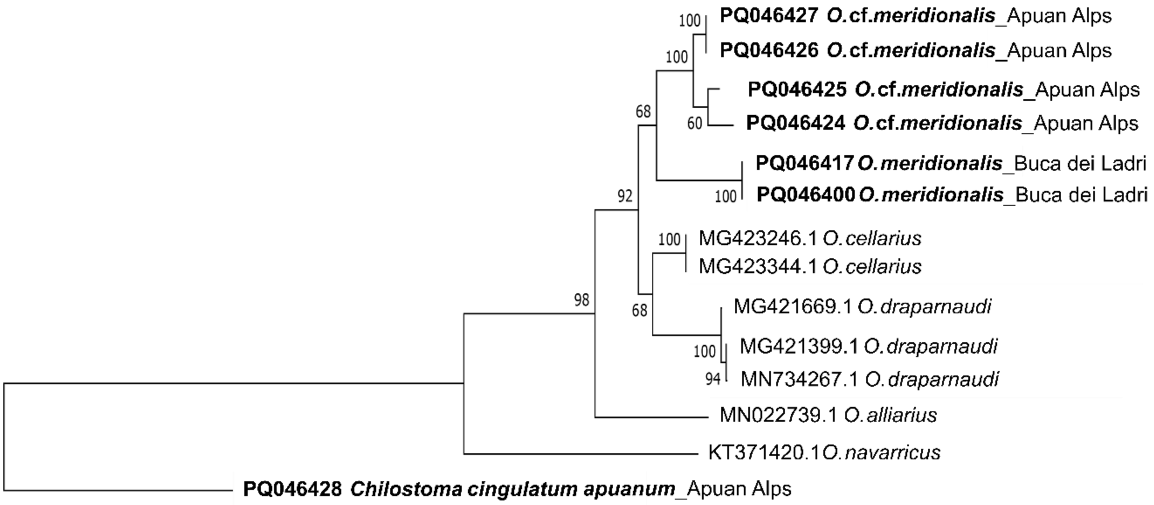


**S9.** Maximum-likelihood tree obtained from dataset consisting of the newly sequences from this study (in bold) and some sequences extracted from GenBank. The numbers represent node support

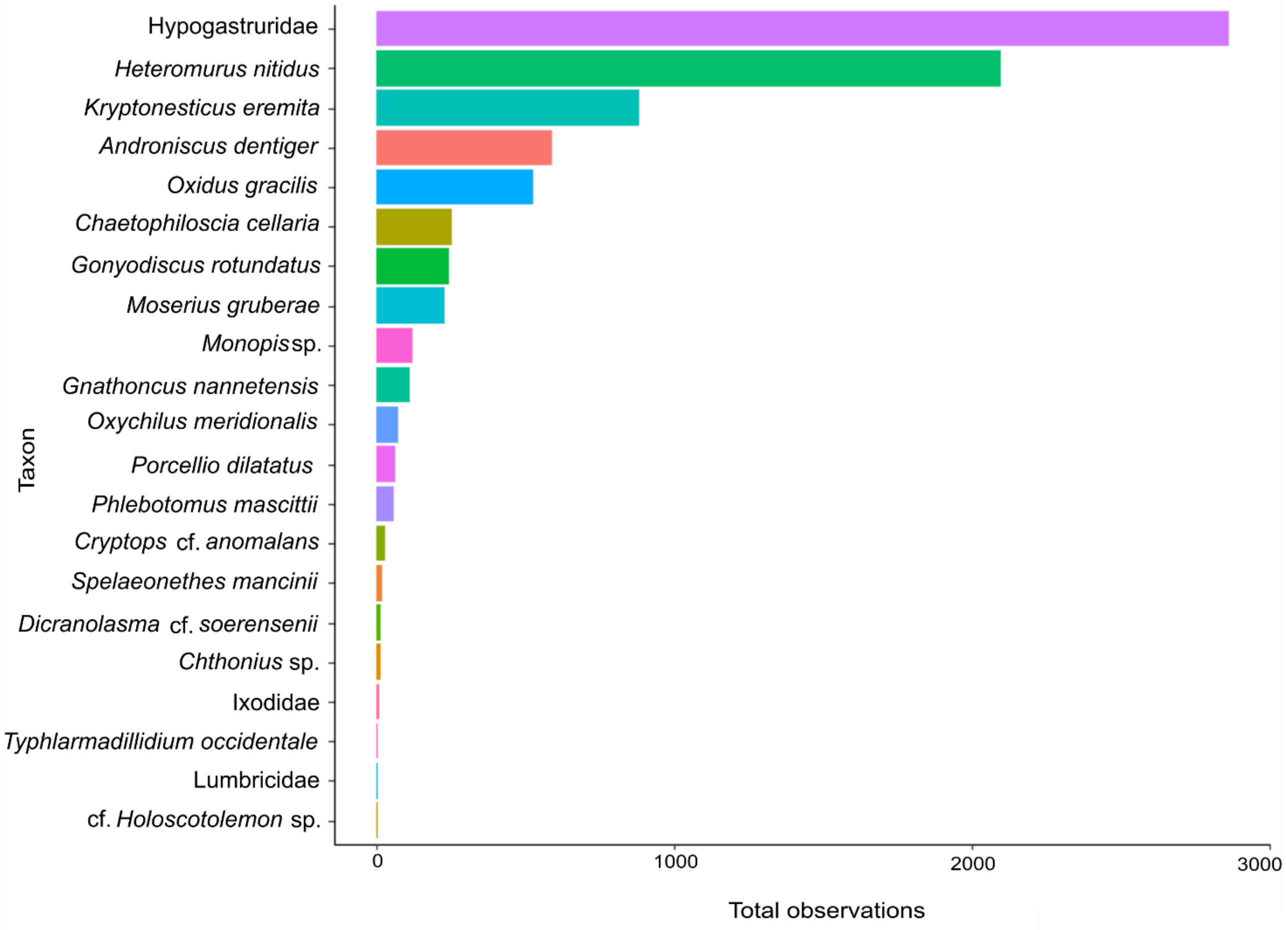


**S10.** Total observations for each taxon detected inside the sampling quadrats

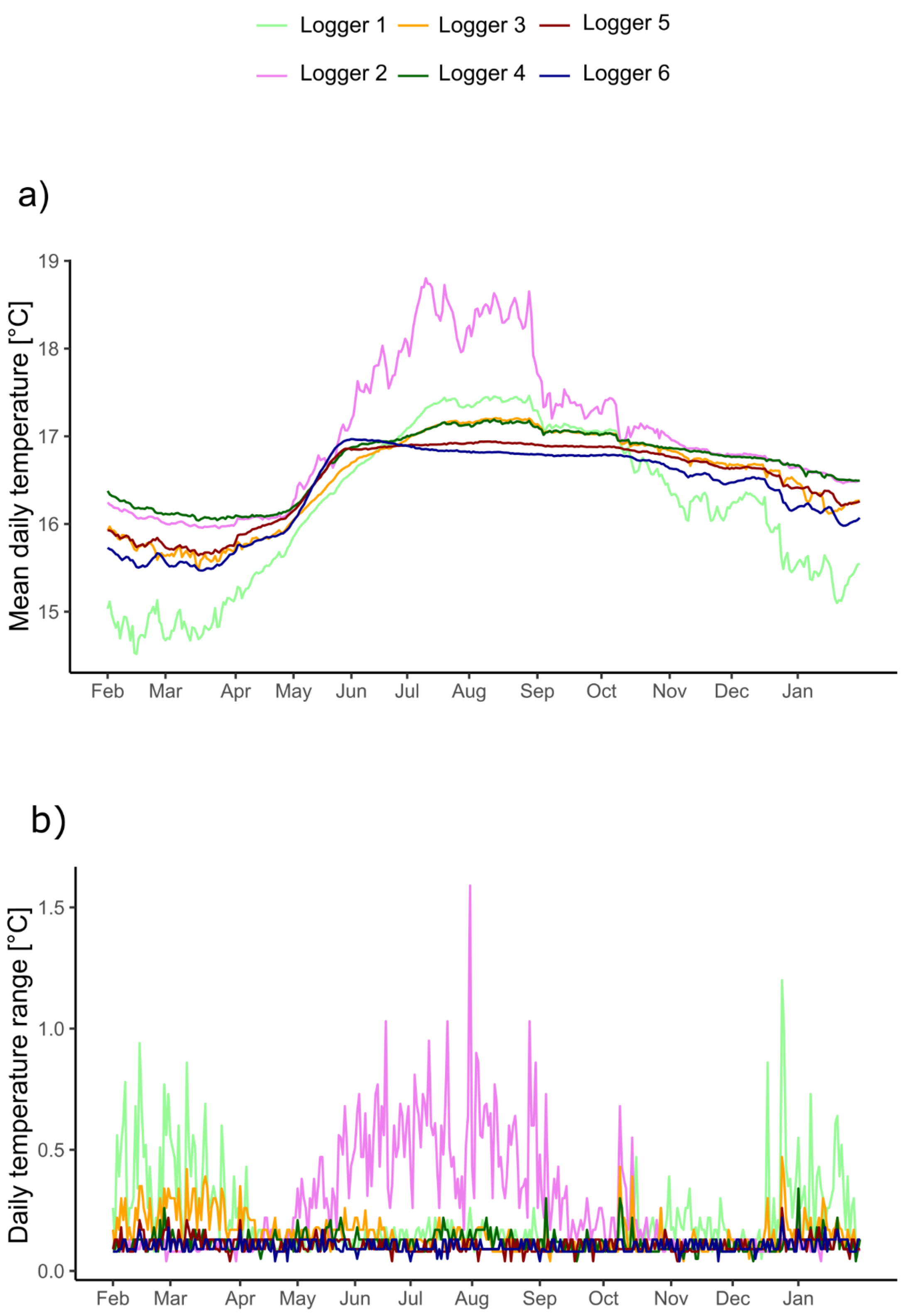


**S11. a**) Annual trend (from February 2022 to January 2023) and **b**) values of daily thermal excursion of the temperature inside the cave. Each curve of different colour represents a datalogger
